# Convergent Skeletal Muscle Cytokine Responses to TFEB Overexpression and Voluntary Wheel Running Reflect Sex-Based Variability in Exercise Adaptations

**DOI:** 10.1101/2025.05.28.655862

**Authors:** Dalton C. Patterson, Allison Birnbaum, Ian Matthews, Constanza J. Cortes

## Abstract

Endurance exercise (running) promotes skeletal muscle remodeling through metabolic and inflammatory signaling cascades. However, the extent to which these responses are sex-dependent remains unclear. Here, we profiled cytokine responses in quadriceps muscle lysates from sedentary, voluntary wheel-running (VWR; 5 weeks), and muscle-specific TFEB-overexpressing (cTFEB;HSACre) male and female mice. Cytokine analysis revealed 40 differentially expressed factors associated with exercise and/or TFEB overexpression, many displaying sex-dimorphic expression patterns. In males, VWR induced significant increases in interleukins (e.g., IL-1α, IL-1β, IL-2, IL-5, IL-17) and chemokines (e.g., MCP-1, CCL5, CXCL9), as well as cytokines involved in TNF signaling (e.g., TNFα, sTNFR1/2, Fas ligand). TFEB overexpression in sedentary males recapitulated many of these cytokine elevations. In contrast, female runner muscle showed limited cytokine activation, with significant changes restricted to IL-3, IL-3Rb, IL-13, and CXCL16. Both sexes exhibited a reduction in IL-4 and an increase in IGFBP-5 with running. Several additional male-specific cytokine profiling responses, including increases in IFNγ, SCF, TPO, VCAM1A, and leptin, further underscored the sex-specificity of exercise-related inflammatory adaptations. These findings demonstrate that skeletal muscle cytokine responses to endurance-like stimuli are profoundly influenced by sex and suggest that male muscle exhibits a broader and/or later remodeling profile than female muscle. Our data also implicate skeletal muscle TFEB-overexpression as a partial molecular mediator of the cytokine shifts observed with exercise, particularly in males, and highlights their potential use as a new prioritization platform for exercise-associated phenotypes.

## INTRODUCTION

Cytokines are a diverse family of intracellular signaling molecules regulating multiple organ system functions and maintenance during aging and disease. Cytokines maintain tissue homeostasis through precise signaling between pro- and anti-inflammatory cytokines, and this balance plays an important role in regulating the body’s response to exercise ^1,2^. There are many sub-families of cytokines, each with unique functions and effects, such as interleukins (ILs), tumor necrosis factor (TNF), interferons (IFNs), chemokines, colony-stimulating factors (CSFs), and growth factors. Exercise modulates the expression and secretion of many of these cytokines in skeletal muscle ^2,3^, where they act as autocrine, endocrine, and paracrine mediators ^4^ contributing to the full metabolic and functional adaptations of long-term exercise ^4^. Indeed, in the context of exercise, the primary source of cytokine responses is the skeletal muscle itself ^1^. This suggests that cytokines play a central signaling role in the musculoskeletal system and beyond. However, the constellation of exercise-activated responses on transcriptional, metabolic, and functional outcomes in skeletal muscle is highly complex ^2^, with the directionality and intensity of these signatures being highly responsive to variations in experimental cohorts, exercise interventions, and measurement timing and methods ^2^, making the isolation of primary effectors from secondary responses difficult ^5^. Overall, most evidence indicates that after an initial pro-inflammatory profile following acute exercise ^2,4^, regular exercise leads to an overall reduction in pro-inflammatory cytokines and an increase in anti-inflammatory cytokine profiles ^3,4,6^, although the degree to which these profiles change remains widely debated. Furthermore, to date, most research into exercise-activated cytokines has remained male-centric, and few studies have examined how sex contributes as a biological variable to exercise-activated cytokine profiles ^2^.

Transcription Factor E-B (TFEB) is a master regulator of mitochondrial metabolism and the endolysosomal network ^7–9^. The expression and function of TFEB are strongly induced in skeletal muscle in response to exercise ^7,10^, and TFEB is required for the full metabolic remodeling observed in skeletal muscle after treadmill training ^7^. We have recently demonstrated that overexpression of TFEB using a pan-skeletal muscle driver (cTFEB;HSACre transgenic mice) is sufficient to drive endurance-like remodeling of quadriceps, gastrocnemius, and soleus muscle in the absence of any exercise activity ^11^. Muscle-TFEB overexpression increased muscle fiber cross-sectional area, decreased the abundance of fast-twitch type IIb fibers, and increased the abundance of slow-twitch type I and IIa fibers in young gastrocnemius muscle ^11^. We also reported increased exercise endurance capacity in treadmill exhaustion tests for young cTFEB;HSACre transgenic mice of both sexes ^11^, and preserved quadriceps and gastrocnemius wet weight (skeletal muscle mass) in aged cTFEB;HSACre transgenic mice relative to their age-matched littermate controls ^11^. Unbiased proteomic profiling of TFEB-overexpressing quadriceps muscles revealed multiple categories associated with cellular metabolism and mitochondrial function, including thermogenesis, oxidative phosphorylation, fatty acid metabolism, and amino acid metabolism ^11^. This is consistent with how acute AAV-mediated TFEB expression in young male skeletal muscle was previously shown to induce the expression of genes involved in mitochondrial biogenesis and oxidative phosphorylation, increasing exercise endurance via improving glycogen metabolism ^7^. We also demonstrated that muscle-TFEB overexpression promotes the secretion of exercise-responsive, muscle-originating factors into circulation (ie. exerkines), including cathepsin B ^11^ and prosaposin ^12^. Importantly, we were also the first to evaluate these local and systemic exercise-like effects across both sexes and identified both sex- and age-dimorphic trajectories associated with TFEB-overexpression in skeletal muscle ^11^. Altogether, this suggests that muscle-TFEB overexpression promotes maintenance of mitochondrial function and bioenergetic reserves, induces running-like fiber type switching, and preserves skeletal muscle function during aging, all profiles reminiscent of those seen after endurance training ^13^. Here, we report that cytokine signatures from cTFEB;HSACre skeletal muscle partially overlap with cytokine signatures from runner muscle, suggesting it represents an intermediate profile between sedentary and exercised states. Furthermore, we find that female runner skeletal muscle cytokine profiles do not follow the same patterns as their male counterparts, but they are reminiscent of female muscle-TFEB overexpressing muscle, highlighting conserved sex-dimorphic responses to exercise and TFEB-mediated physiology. Altogether, we present evidence highlighting the potential of cTFEB;HSACre transgenic mice as a powerful orthogonal tool for examining and prioritizing skeletal muscle exercise-associated profiles.

## RESULTS

### Running-activated cytokine profiles are sex-dimorphic

To profile the shared and unique cytokine responses associated with endurance-like muscle remodeling in male and female mice, we generated quadriceps muscle protein lysates from young (4.5-months-old) sedentary, runners (voluntary wheel running, VWR) or cTFEB;HSACre transgenic mice of both sexes (n=3-4/group) (**Figure 1A**). Mice in the running group were provided 24-hour access to an in-cage running wheel for 5 weeks, whereas the sedentary group remained in cages with a locked wheel. This length of running intervention has been previously shown to be sufficient to promote exercise-associated muscle adaptations, including increased mitochondrial activity ^14,15^, increased capillary density ^16^, and fiber type switching ^16,17^ in young mice. We chose this voluntary running paradigm to avoid stress-induced activation of inflammation associated with long-term forced treadmill running ^18,19^. Consistent with previous reports monitoring similar durations of voluntary running in wild-type mice at similar ages ^20^, we observed continuous running behavior across 5 weeks for both male and female mice (**Figure 1B**). Interestingly, female mice displayed a higher preference for running over their male littermate counterparts during the wheel acclimation period (**Figure 1B**, week 1). Male runners caught up to female runners in both the average (not shown) and cumulative wheel counts by week 3 and remained comparable till the end of the experiment (**Figure 1B**, weeks 2-5). Indeed, males and females had completed similar total wheel counts by the end of the study (**Figure 1C**). Body weight measurements at the end of the running trials confirmed that 5 weeks of VWR reduced total body weight in male (and trending for female) runners (**Figure 1D**), consistent with the well-known benefits of VWR on fat/lean mass ^20^. As we have shown previously ^11^, no differences in body weight were observed in young male/female cTFEB;HSACre transgenic mice when compared to their littermate controls (**Figure 1D**).

**Figure 1:**
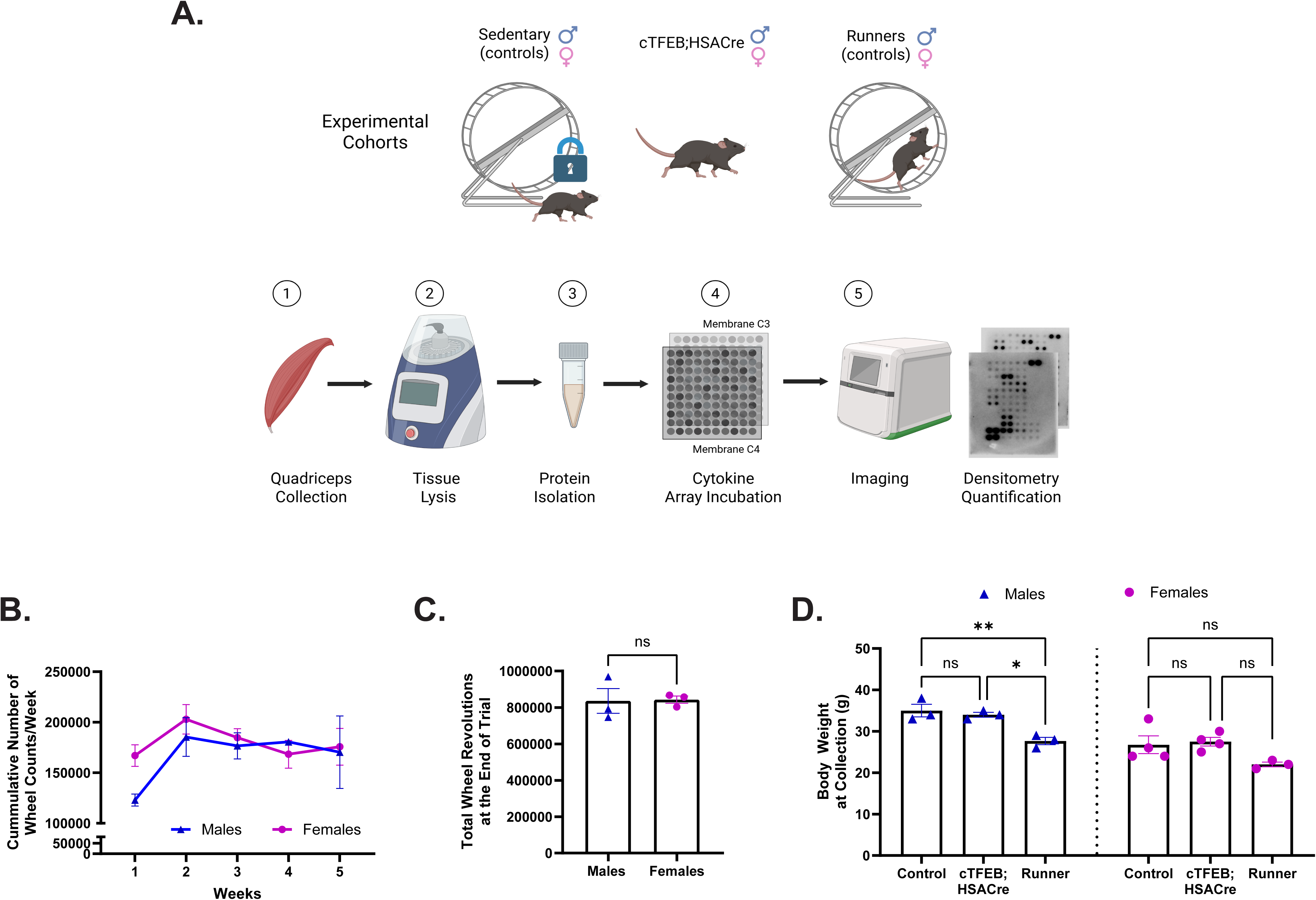
Voluntary Running Wheel (VRW) Performance and Body Weight in Experimental Cohorts. **(A)** Design schematic highlighting experimental cohorts: sedentary (n=4 M/F), cTFEB;HSACre transgenic (n=4 M/F) and runner (n=3 M/F) mice, 4.5 months of age at the beginning of the experiment. Sample processing and analysis pipeline is shown below. **(B)** No significant differences in cumulative wheel counts/week throughout the course of the experiment or, **(C)** total wheel counts at the end of the experiment for both sexes. **(D)** Significantly reduced body weight in male runners (and trending for female runners) at collection, no differences in cTFEB;HSACre transgenic mice body weight relative to sedentary controls. N.s: non-significant, * P ≤ 0.05, ** P ≤ 0.01. Data is represented as mean ± SEM.

Next, we simultaneously assessed 96 mouse inflammation and immune system responsive-cytokines in quadriceps muscle total protein lysates from all groups (**Figure 1A**). Of these, we identified a total of 40 differentially expressed cytokines, some of which were uniquely associated with running (17), with muscle-TFEB overexpression (4), or following similar directional changes under both conditions (19). Notably, most of these also exhibited a sex-dimorphic pattern, with robust exercise and cTFEB;HSACre signatures in males that were absent in females. We describe our findings below.

#### Interleukins (ILs)

Interleukins (ILs) are a group of proteins involved in communication with leukocytes by binding to specific receptors. ILs are mainly synthesized by monocytes, macrophages, T cells, and endothelial cells, but evidence suggests myofibers and myoblasts can also produce ILs ^21^. ILs are highly responsive to both the timing and intensity of the exercise bout, with differential effects on the pro-inflammatory and anti-inflammatory branches of the family ^3,6,21^. In our hands, and similar to what has been previously shown ^21^, we detected significant increases in pro-inflammatory cytokines IL-1α, IL-1β, IL-2, IL-3, IL-5, IL-12 p40/70, IL-12 P70 (not shown), and IL-17 in skeletal muscle lysates from male young runner mice (**Figure 2**). We did not detect any changes in IL-6 levels in runner muscle lysates of either sex (**Figure 2H**), consistent with its role as an early responder to acute bouts of endurance exercise. Interestingly, although many of these cytokines appeared to trend in the same direction as their male counterparts, only IL-3 was also significantly elevated in the muscle of female runners (**Figure 2D**). Furthermore, pro-inflammatory cytokine IL-3Rb (**Figure 2E**) was found to be significantly increased in female (but not male) muscle lysates after 6 weeks of voluntary wheel running. Running also robustly decreased the abundance of anti-inflammatory cytokine IL-4 in both sexes (**Figure 2F**), and increased levels of anti-inflammatory IL-10 (males only) (**Figure 2I**) and IL-13 (females only) (**Figure 2K**). Interestingly, out of these, both IL-5 (**Figure 2G**) and IL-17 (**Figure 2L**) were also significantly upregulated in the muscle lysates of age-matched male (but not female) cTFEB;HSACre transgenic mice, even in the absence of running.

**Figure 2:**
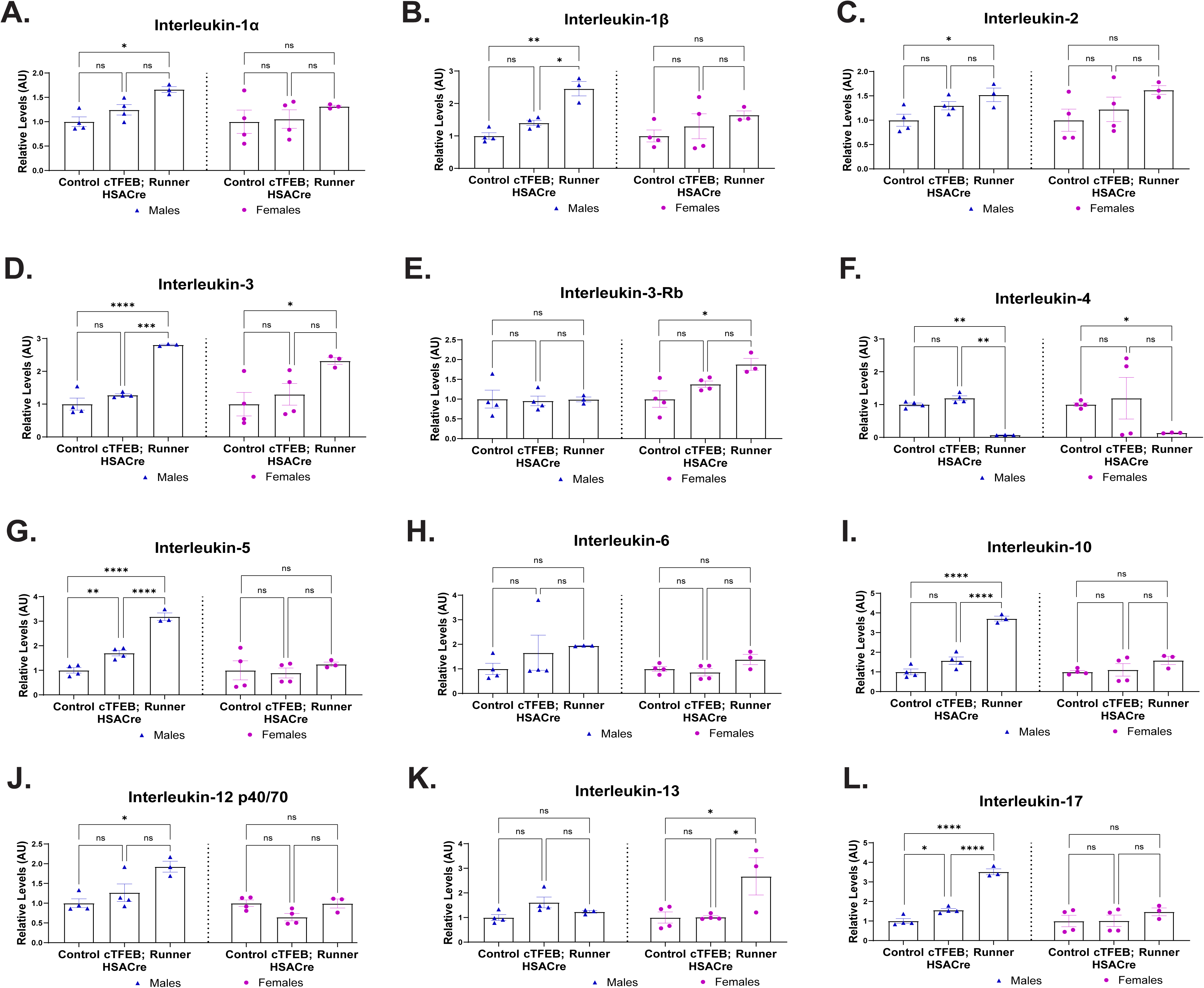
Interleukin cytokine profiles of cTFEB;HSACre and runner skeletal muscle lysates. **(A-L)** Densitometry analysis of interleukin abundance in skeletal muscle protein lysates of all cohorts normalized to respective sedentary baseline for each sex. Arbitrary units. * P < 0.05, ** P < 0.01, *** P < 0.001, **** P < 0.0001. Data is represented as mean ± SEM.

#### Chemokines

Chemokines are another large family of exercise-responsive cytokines ^1,3,21^, which function to stimulate the recruitment of leukocytes and other immune cells from circulation into solid tissues (chemotaxis). They can be divided into two families, with conserved sequential (CC) or proximal (CXC) cysteine residues. Examination using our ELISA approach revealed nine differentially expressed chemokines in skeletal muscle lysates from male young runner mice (**Figure 3**), including MCP-1/CCL2, MCP-5, RANTES/CCL5, Eotaxin 1/CCL11, TARC/CCL17, MIP-3 alpha/CCL20 and MIG/CXCL9. Similar to what we observed in our interleukin panel, these chemokine levels did not change in young female runner skeletal muscle relative to their baseline sedentary controls (**Figure 3**), although we detected an additional significant increase in CXCL16 levels in female (but not male) runner muscle (**Figure 3K**). TFEB-overexpressing muscle also displayed significant increases in MCP-1 (CCL2) (**Figure 3A**) and TARC (CCL17) (**Figure 3F**) to similar levels as their runner littermates, and CXCL16 to a level in between sedentary and runner littermates (**Figure 3K**), following the same sex-specific patterns that we saw in runner chemokine profiles (**Figure 3**). TFEB-overexpressing muscle lysates displayed one unique significant increase in CXCL family member SDF-1 alpha/CXCL12 relative to their non-transgenic sedentary control littermates (**Figure 3J**).

**Figure 3:**
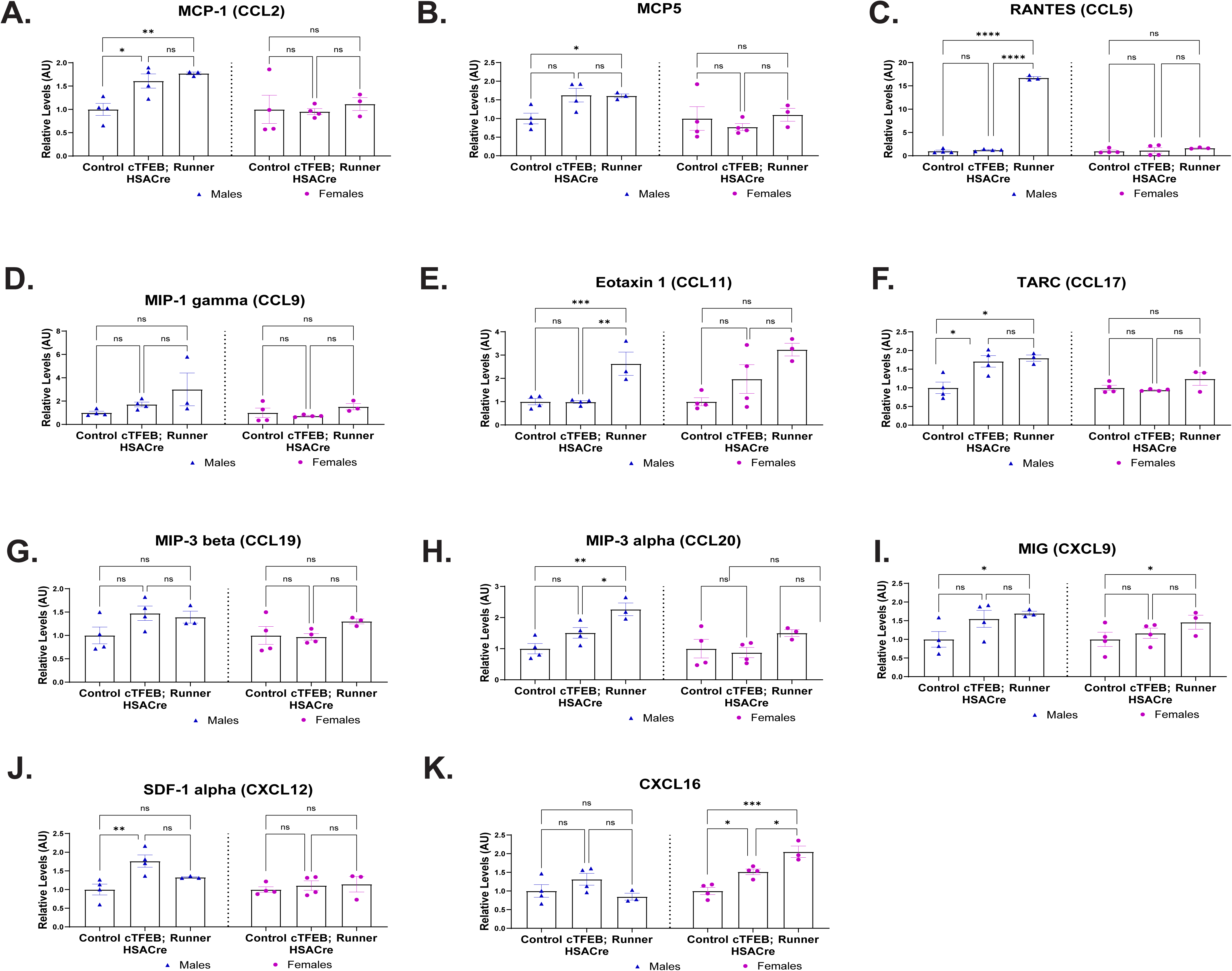
Chemokine cytokine profiles of cTFEB;HSACre skeletal muscle lysates. **(A-K)** Densitometry analysis of chemokine abundance in skeletal muscle protein lysates of all cohorts normalized to respective sedentary baseline for each sex. Arbitrary units. * P < 0.05, ** P < 0.01, *** P < 0.001, **** P < 0.0001. Data is represented as mean ± SEM.

#### Tumor Necrosis Factor (TNF) Signaling

Tumor necrosis factor alpha (TNF-α) is a pluripotent cytokine generally produced by macrophages and adipocytes and is a major mediator of the acute inflammatory response ^22^. TNF-α has two membrane-bound receptors that can be cleaved into soluble isoforms involved in its signaling cascade and likely modulate its dual effects on inflammation: sTNFR1 is constitutively expressed on most cell types, and its signaling is mainly pro-inflammatory, whereas sTNFR2 is primarily restricted to subsets of immune cells, glial cells, and cardiomyocytes, and its signaling can be anti-inflammatory and promote cell proliferation instead ^22^. TNF levels increase with endurance exercise, consistent with what we found in this study (**Figure 4**). Male runner muscle had significantly elevated levels of cytokines involved in the TNF signaling axis, including TNFα, soluble TNFR1, soluble TNFR2, CD40/TNFRSF5, and Fas ligand (**Figure 4A-E**). We found that male cTFEB;HSACre muscle also displayed significant elevations in these cytokines compared to their sedentary littermate controls, to a degree indistinguishable from runner muscle (**Figure 4A-E**). Furthermore, cTFEB;HSACre muscle also had significantly higher cytokine CD30 /TNFRSF8 levels (**Figure 4F**), another member of the TNF signaling axis. Once again, no changes were detected for any TNF-associated cytokines in the skeletal muscle of female cTFEB;HSACre or runner mice (**Figure 4**).

**Figure 4:**
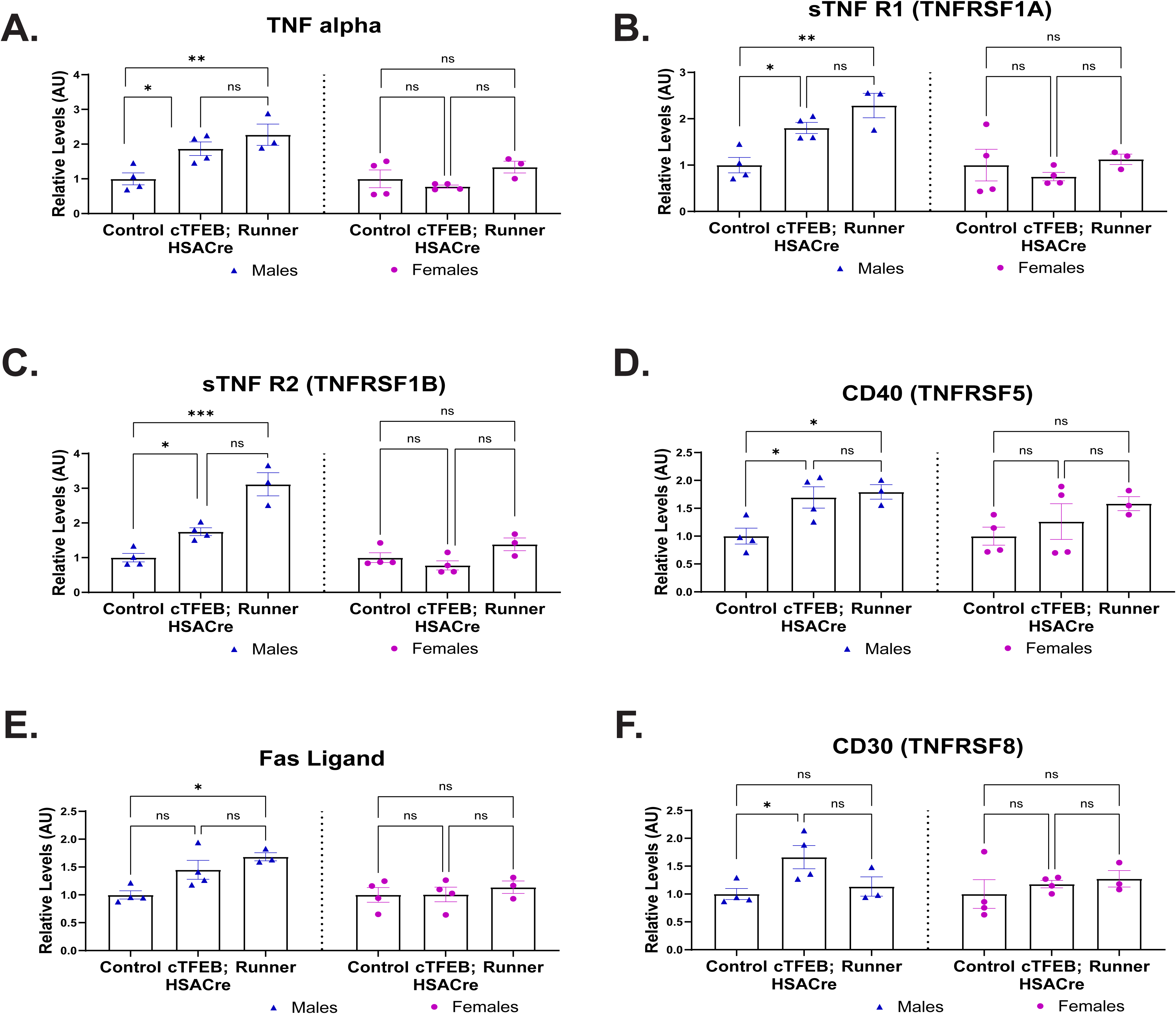
TNF signaling cytokine profiles of cTFEB;HSACre skeletal muscle lysates. **(A-F)** Densitometry analysis of TNF cytokine family abundance in skeletal muscle protein lysates of all cohorts normalized to respective sedentary baseline for each sex. Arbitrary units. * P < 0.05, ** P < 0.01, *** P < 0.001. Data is represented as mean ± SEM.

#### Additional Cytokines

Interferon-gamma (IFNγ) increases in human plasma after regular moderate exercise ^23^ and immediately after acute moderate exercise ^24^ and has been linked to the anti-inflammatory benefits of regular physical activity. We detected a significant increase in IFNγ abundance in muscle lysates from our runners and our cTFEB;HSACre transgenic muscle, although to a lesser degree (**Figure 5A**). No changes were detected for IFNγ levels in the muscle of female mice (**Figure 5A**).

**Figure 5:**
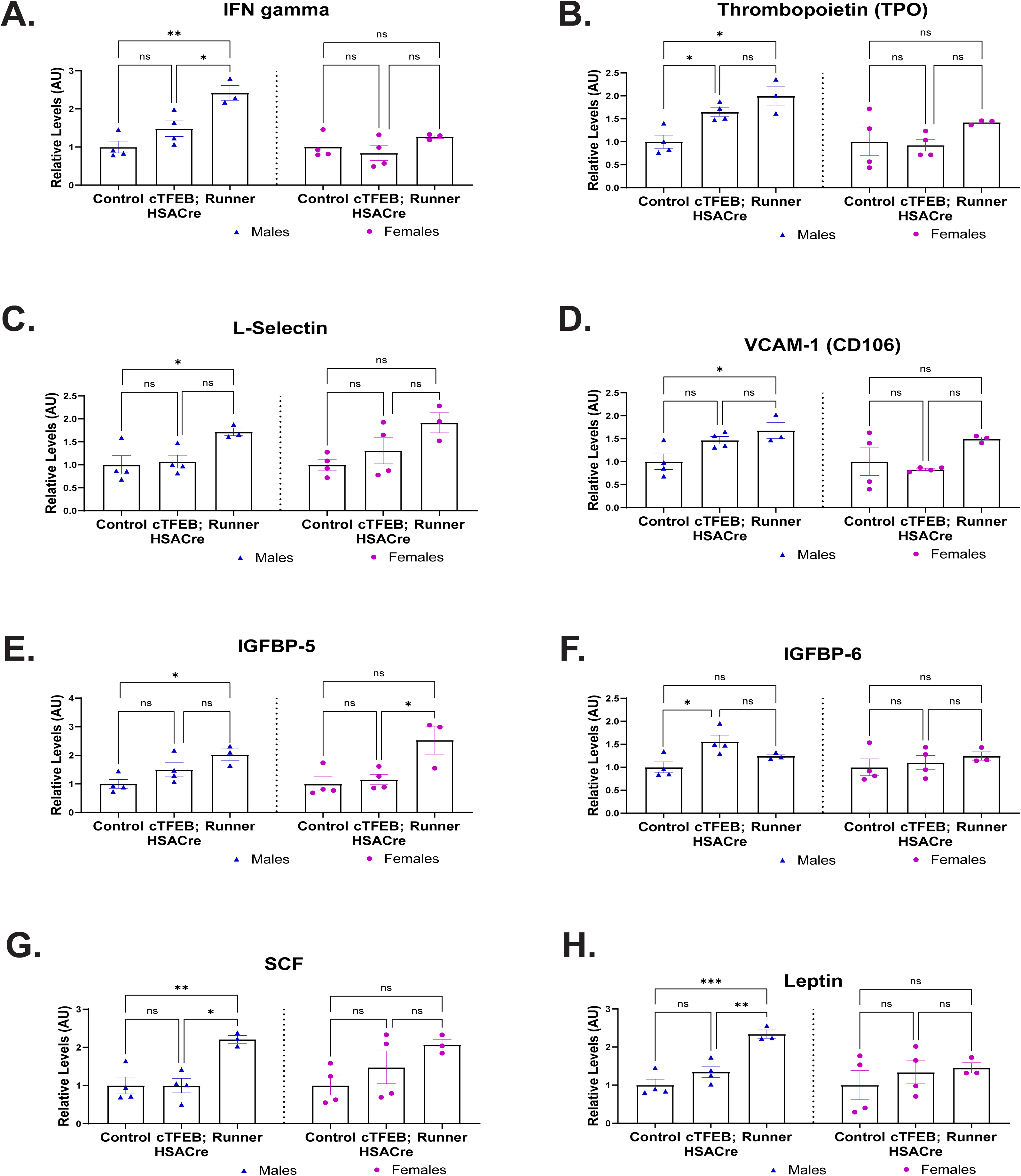
Orphan signaling cytokine profiles of cTFEB;HSACre skeletal muscle lysates. **(A-H)** Densitometry analysis of additional cytokine family abundance in skeletal muscle protein lysates of all cohorts normalized to respective sedentary baseline for each sex. Arbitrary units. * P < 0.05, ** P < 0.01, *** P < 0.001. Data is represented as mean ± SEM.

Thrombopoietin (TPO), a glycoprotein hormone that regulates platelet production and may improve osteogenesis in the absence of mechanical loading ^25^, was also significantly elevated in both male runner and male cTFEB;HSACre muscle (**Figure 5B**). L-selectin and Vascular cell adhesion molecule 1 A (VCAM1A) are cell adhesion molecules that play a role in inflammation responses and leukocyte adhesion, potentially modulating immune cell infiltration during inflammation. We found increases in both of these cytokines in the muscle of our male (and trending for female) runners (**Figure 5C-D**).

IGFBP-5, an insulin-like growth factor binding protein that stimulates IGF-1 signaling, plays a complex role in skeletal muscle, influencing both muscle growth and atrophy ^26^, and appears to decrease in the interstitial muscle space in human muscle after acute exercise ^27^. Importantly, IGFBPs are known to have IGF-independent actions in cell migration, cell growth, and apoptosis ^28^. We found a significant increase in IGFBP-5 levels in both male and female runner muscle relative to their sedentary littermate controls (**Figure 5E**). IGFBP-6, another insulin-like growth factor binding protein family member that inhibits IGF-1 signaling, is increased in the muscle of mice selected for high running performance ^29^ and in regenerating skeletal muscle ^30^. We found a significant increase in abundance levels of IGFBP-6 in male cTFEB;HSACre (but not runner) muscle (**Figure 5F**).

Stem Cell Factor (SCF) is a cytokine that recruits stem cells from the bone marrow into sites of injury via activation of receptor tyrosine kinase c-Kit ^31^. SCF was also found to be significantly elevated in male runner (but not cTFEB;HSACre) muscle, with similar trends in female runner muscle (**Figure 5G**). Finally, we also observed significantly increased leptin levels, a hormone commonly synthesized by fat tissue, which has been noted to regulate energy balance and metabolism ^32^, in male runner muscle lysates (**Figure 5H**).

## DISCUSSION

Exercise alters cytokine production in skeletal muscle ^1–3,21^, but the role biological sex plays in these cytokine profiles remains largely unknown. Here, we provide direct evidence demonstrating that female skeletal muscle cytokine responses to 5 weeks of VWR significantly differ from their male counterparts, and that our muscle-TFEB overexpression mimics aspects of this sex-dimorphic cytokine response to voluntary wheel running.

Most exercise-regulated cytokines examined here play myriad roles in the immune and non-immune systems ^3^. Indeed, pro- and anti-inflammatory cytokines are altered by acute and chronic exercise in rodents and humans ^33^, although their physiological effects are context-, timing-, and cell-type specific ^4^. It has been suggested that the local beneficial impact of chronic exercise is mediated, at least in part, by exercise-induced ‘hormesis’ ^34^, whereby a low level of stress (in this case, associated with the increased metabolic demand of the running bout) upregulates existing cellular and molecular pathways that improve physiological hallmarks ^4^. Thus, specific combinations of cytokines (sometimes with opposing functions) and the directionality, magnitude, and duration of these changes likely underlie metabolic and architectural remodeling of skeletal muscle in response to running. Consistent with this hypothesis, we identified multiple inflammation-associated cytokines (ILs, TNF signaling, chemokines) that are significantly elevated in male skeletal muscle after 5 weeks of VWR. Many of these have been reported to be exercise-responsive before, both within skeletal muscle ^35^ and in circulation ^21,35^. Interestingly, one week of VWR has been shown to induce more cytokine changes than either an acute low-intensity or high-intensity exercise in mice ^35^, further highlighting the importance of timing and intervention vs. recovery phases during exercise studies. The fact that pro-inflammatory cytokines remain elevated in our 5-week VWR intervention suggests that these cytokine profiles may be associated with ongoing exercise-dependent muscle remodeling rather than muscle damage, although this hypothesis remains to be tested.

Many of the differentially expressed cytokines we identified in this study are chemoattractants (chemokines), adhesion molecules that facilitate extravasation (VCAM1, L-Selectin), or immune-system signaling molecules (interleukins). Skeletal muscle has both resident and exercise-responsive immunocyte populations, including macrophages, T-regs, and neutrophils, and exercise has well-known effects on immune cell circulationand tissue-resident populations ^3,4,33^. Importantly, the fold changes we observed in most canonically known pro-inflammatory interleukins (IL-1α, IL-1β, IL-2, IL-5, IL-12 p40/70, and IL-17) are mild (between 2-4-fold over sedentary controls), and may reflect tissue-remodeling responses or chronic sub-pathological levels of inflammation (’inflammation hormesis’). Additionally, the chemokine profiles we detected suggest a local increase in the production of chemoattractants targeting monocytes (MCP-1/CCL2, MCP-5), macrophages (MIP-1 gamma/CCL9), T-cells (MIP-3 beta/CCL19, MIP-3 alpha/CCL20, MIG/CXCL9 and TARC/CCL17), eosinophils (Eotaxin 1/CCL11) and leukocytes (RANTES/CCL5). It is currently unclear whether these profiles reflect an increase in local immunocyte signaling or an exercise-dependent infiltration of new immune cells into runner and/or cTFEB;HSACre muscle. Although we cannot rule out this possibility, the type and duration of exercise the mice were subjected to in our study generally do not cause significant muscle damage ^35^, suggesting that if these are new infiltrating immunocytes, they may be involved in tissue remodeling rather than tissue repair responses.

We ^11^, and others ^7,11^, have previously shown that muscle-TFEB overexpression engages multiple exercise-like effects in skeletal muscle, including mitochondrial remodeling, fiber type switching, and expansion of the endolysosomal network. Unbiased proteomics analysis identified many classical exercise-activated networks in TFEB-overexpressing quadriceps muscle, including proteostasis, mitochondrial metabolism, fatty acid metabolism, and amino acid metabolism ^11^. Furthermore, we also confirmed that muscle-TFEB overexpression increases the secretion of exercise-responsive factors into circulation ^11^, which promotes exercise-like remodeling in the Central Nervous System ^11,12^. Here, we expand our original analysis of the effects of TFEB-overexpression in skeletal muscle and demonstrate that it also mimics cytokine profiles associated with voluntary wheel running. Indeed, we detected significantly increased levels of interleukins (IL-5, IL-17), chemokines (MCP-1/CCL2, TARC/CCL-17, CXCL16), and TNF signaling (TNFα, sTNFR1, sTNFRII, CD40/TNFRSF5) in cTFEB;HSACre skeletal muscle to levels similar (or intermediate) to those seen in runner muscle.

Interestingly, the strongest concordance in cytokine profiles between cTFEB;HSACre and runner muscle was observed in the TNF signaling pathway. Although commonly associated with muscle tissue loss in conditions of cachexia or sarcopenia, TNF-α also has important roles in muscle repair ^36^ and growth and differentiation ^37,38^, and can be produced locally by myocytes and myoblasts ^21,39^. The increases we observed in TNF signaling cytokines in either cTFEB;HSACre or runner lysates are much lower than those reported for muscle pathological conditions, suggesting that they may instead represent immunomodulatory states associated with these anabolic functions of TNF signaling. In agreement with this, in cTFEB;HSACre muscle lysates we observed significant increases in sTNFRII, which is considered mostly anti-inflammatory and promotes cell proliferation ^22^. Interestingly, TNFα also induces the surface expression of various adhesion molecules to promote the migration of leukocytes at sites of inflammation (including vascular cell adhesion molecule-1 (VCAM-1) and L/E-selectin), which we also found were significantly increased in runner muscle lysates. As before, whether these increases reflect local production of TNF-signaling cytokines or an increased infiltration of immunocytes into either cTFEB;HSACre or runner muscle remains unknown.

One of our most unexpected findings was the profound sex bias exhibited by almost all the cytokines in this panel. Out of 40 differentially abundant cytokines profiled in our study, only two followed the same patterns in male and female runner muscle (IL-3 and IL-4). Furthermore, three cytokines appeared to be female-specific (IL-3Rb, IL-13 and CXCL16), with significant increases in female (but not male) runner muscle lysates. It is important to note that we have confirmed human TFEB transgenic protein expression levels to be 3-fold higher in male than female cTFEB;HSACre skeletal muscle ^11^. While this may account for some of the sex-specific changes in cytokines we report here, the fact that the cytokine expression pattern of female runner-muscle was similar to what we observed in female cTFEB;HSACre muscle instead suggests that conserved sex-dimorphic signals may control cytokine abundance across both models.

Sex differences in exercise responses remain a profoundly understudied topic of research ^2,40^, and in general, we struggled to identify previous literature describing female-specific changes to many of these cytokines in response to any exercise modality. This renders interpretation of these female-specific patterns (or lack thereof) difficult, but we believe a set of overlapping and/or non-mutually exclusive scenarios could be at play here. In our hands, both sexes displayed similar running patterns (total wheel revolutions at the end of the 5 weeks of VWR), and similar trends towards decreases in body weight between our sedentary and runner groups of both sexes. This suggests that the intensity of the running intervention had similar systemic effects on the fat/lean mass of our groups, although specific biological responses in adipose tissue, liver, and other highly metabolic tissues were not examined in this work. Another option is the timing of the cytokine responses: our analysis was performed at the end of 5 weeks of running. We did observe a significantly higher number of cumulative wheel counts during Weeks 1-2 in our female runners relative to their male littermates. This may indicate female runners may reach peak cytokine increases earlier than their male counterparts, with compensatory responses returning these responses to baseline by the time we collected our samples. Finally, it could also be that female and male skeletal muscle possess different steady-state profiles for these cytokines, which makes exercise or muscle-TFEB-activated changes different across sexes, although we did not examine this possibility in this current work.

Both sexes display the well-known benefits of exercise on skeletal muscle, suggesting converging metabolic and functional outcomes despite differences in timing, abundance, or cross-talk between these immune-associated cytokines. Thus, it may be that a specific combination of cytokines (with diverse and opposing functions), whose direction, magnitude, and duration of change, which also appear to be highly sex-dimorphic, underlie exercise-induced hormesis. In agreement with this, the recently released Molecular Transducers of Physical Activity Consortium (MoTrPAC) dataset has also highlighted the diverse temporal and sex-dynamics of multi-omic responses to endurance training in young rats ^2^. Targeted interrogation of the MoTrPAC data hub for transcriptional and proteomic levels of detected cytokines also present in the cytokine array utilized in this study reveals trends towards sex-dimorphic responses for IL-1α, IL-2, IL-12 p40/70, IL-13, and IFN-γ.

Furthermore, the temporal responses of IL-1α and IL12, for example, are also increased during week 1 and disappear by week 4 of the treadmill training in the gastrocnemius of female runner rats ^2^, consistent with our hypothesis that biological sex may play an important role in exercise-activated cytokine timing and abundance. Together, these studies underscore the fundamental need to examine sex differences that contribute to exercise physiology.

Cytokine networks and their responses to exercise signaling must be tightly regulated to limit damage whilst maintaining the benefits, some of which appear to be recapitulated in TFEB-overexpressing muscle. Overall, the similarities we observed on our cytokine profiling of runner and cTFEB;HSACre skeletal muscle suggest that muscle-TFEB overexpression may prime skeletal muscle to an intermediate exercise-like state. Indeed, we see cTFEB;HSACre skeletal muscle adopting both characteristics of the initial phase of the immune response during exercise ^4^ (with pro-inflammatory states characterized by cytokines such as IL-5, IL-17 and TNFα) but also of the later phases of the exercise-activated immune response (with a preponderance of anti-inflammatory states, characterized by IL-10, IL-13 and sTNFR2) ^4^. Furthermore, for several chemokines that did not reach statistical significance (MCP-5, MIP-3 alpha, MIG, and leptin, for example), muscle-TFEB overexpressing samples displayed clear trends towards similar increases as those seen in our runner lysates, suggesting additional nodes of shared signaling between these two models. This highlights the potential of the cTFEB;HSACre transgenic mouse model as a new orthogonal prioritization platform to identify and validate exercise-associated signaling, including cytokines, secreted factors ^7,10^, and age-associated geroprotective benefits ^11,12^.

## METHODS

### Animals

We have previously described the generation of fxSTOP-TFEB transgenic and cTFEB;HSACre double transgenic mice ^7^. For this work, we utilized controls or cTFEB;HSACre double transgenic male and female littermates (∼4-5 months of age). In our hands, we have not detected any differences in muscle phenotypes on fxSTOP-TFEB+ or HSA-Cre+ single transgenic animals compared to wild-type littermates ^7^, so they are all combined into our ‘control’ groups. Control animals were randomly assigned to each sedentary/runner group (see below). Body weight was measured prior to euthanasia and tissue collection and was accidentally not collected for one male control and one male cTFEB;HSACre animal (see Figure 1D). All animals are in the C57BL/6J genetic background, and we utilized similar numbers of males and females. All animal experimentation adhered to NIH guidelines and was performed by blinded investigators (when possible), and was approved by and performed in accordance with, the University of Alabama at Birmingham (UAB) and the University of Southern California (USC) Institutional Animal Care and Use.

### Exercise Protocol

Four-month-old control mice were singly housed with access to in-cage locked (sedentary) or free (running) wheels for 5 weeks at the UAB Small Animal Phenotyping Core. To equilibrate the influence of single housing on animal behavior, each mouse from sedentary/runner cohorts was single caged in standard vivarium polypropylene cages (290L×L180L×L160 mm) throughout the experiment in an incubator that maintained a constant temperature. Each running mouse had free access to a running wheel (13 cm in diameter) connected to a counter to record the running distance (Lafayette Science’s AWM activity monitoring software). This in-cage wheel running system for mice collects the number of revolutions and the timing of the revolutions for each cage every 10 minutes. Sedentary controls were also given free access to a locked wheel to control for enrichment. Investigators monitored animal welfare and physical activity daily.

### Tissue collection

Animals were anesthetized with 3.8% Avertin Solution prior to tissue collection. All animals received a transcardial perfusion with 60 mL of ice-cold 1x PBS. Quadriceps skeletal muscle tissue was flash-frozen in liquid nitrogen for protein extraction and downstream analyses.

### Protein Extraction

Protein lysates from muscle tissue were prepared following standard protocols with minor modifications ^11,41,42^. In short, 50 mg of flash frozen perfused quadriceps muscle were placed in plastic tubes with silica beads (MP Biomedical,116913100) and homogenized in the FastPrep-24 5G bead grinder and lysis system (MP Biomedical, 116005500) in RIPA Lysis and Extraction Buffer (Invitrogen, 89900), 1X Halt Protease and Phosphatase Inhibitor Cocktail (Invitrogen, 78442), and 1% SDS. After standard centrifugation steps, protein concentration was quantified using a Pierce BCA Protein Assay (23227).

### Cytokine Array

Cytokine antibody arrays (Abcam, ab193659) were used according to the manufacturer’s protocol. For skeletal muscle analysis, 50 µg of previously prepared quadriceps protein lysate was incubated with the membranes precoated with captured antibodies (C3: 62 targets and membrane C4: 34 targets). Targets analyzed were as follows: Axl, Bfgf, BLC (CXCL13), CD30 Ligand (TNFSF8), CD30 (TNFRSF8), CD40 (TNFRSF5), CRG-2, CTACK (CCL27), CXCL16, CD26 (DPPIV), Dtk, Eotaxin-1 (CCL11), Eotaxin-2 (MPIF-2/CCL24), E-Selectin, Fas Ligand (TNFSF6), Fc gamma RIIB (CD32b), Flt-3 Ligand, Fractalkine (CX3CL1), GCSF, GITR (TNFRSF18), GM-CSF, HGFR, ICAM-1 (CD54) IFN-gamma, IGFBP-2, IGFBP-3, IGFBP-5, IGFBP-6, IGF-1, IGF-2, IL-1 beta (IL-1 F2), IL-10, IL-12 p40/p70, IL-12 p70, IL-13, IL-15, IL-17A, IL-17 RB, IL-1 alpha (IL-1 F1), IL-2, IL-3, IL-3 R beta, IL-4, IL-5, IL-6, IL-7, IL-9, I-TAC (CXCL11), KC (CXCL1), Leptin, Leptin R, LIX, L-Selectin (CD62L), Lungkine (CXCL15), Lymphotactin (XCL1), MCP-1 (CCL2), MCP-5, M-CSF, MDC (CCL22), MIG (CXCL9), MIP-1 alpha (CCL3), MIP-1 gamma, MIP-2, MIP-3 beta (CCL19), MIP-3 alpha (CCL20), MMP-2, MMP-3, Osteopontin (SPP1), Osteoprotegerin (TNFRSF11B), Platelet Factor 4 (CXCL4), Pro-MMP-9, P-Selectin, RANTES (CCL5), Resistin, SCF, SDF-1 alpha (CXCL12 alpha), Sonic Hedgehog N-Terminal (Shh-N), TNF RI(TNFRSF1A), TNF RII (TNFRSF1B), TARC (CCL17), I-309 (TCA-3/CCL1), TECK (CCL25), TCK-1 (CXCL7), TIMP-1, TIMP-2, TNF alpha, Thrombopoietin (TPO), TRANCE (TNFSF11), TROY(TNFRSF19), TSLP, VCAM-1 (CD106), VEGF-A, VEGFR1, VEGFR2, VEGFR3, and VEGF-D. After washing, the membranes were incubated with the provided streptavidin-HRP secondary detection antibodies. Membranes were developed using the Chemi Reagent Mix kit. Array blot images were captured and visualized using the ChemiDoc MP Imaging System (Bio-Rad), and the intensity of each spot (densitometry) in the captured images was analyzed using the publicly available software ImageJ as per kit instructions. To determine fold changes associated with cTFEB;HSACre or running status, relative cytokine levels were all normalized to sedentary control values for each sex. Results are expressed as arbitrary units/fold change over sedentary controls for each sex. Only cytokines with significant differences are shown.

### Statistical Analysis

All data were analyzed by one-way ANOVA within sex with post hoc comparisons using GraphPad Prism 10.4.2 (La Jolla, CA) and are represented as means and standard error of the means. All data analyses were conducted in a blinded fashion. All data were prepared for analysis with standard spreadsheet software (Microsoft Excel).

## ACKNOWLEDGEMENTS

We thank members of the lab, past and present, as well as former and current colleagues and collaborators, for their helpful contributions.

## AUTHOR CONTRIBUTIONS

D.C.P., A.B., and I.M., performed the experimental work and reviewed the manuscript. D.C.P and C.J.C. wrote and edited the manuscript. C.J.C. conceptualized the experiments outlined here and obtained (or helped obtain) all associated funding.

